# Gene disruption by structural mutations drives selection in US rice breeding over the last century

**DOI:** 10.1101/2020.08.26.268342

**Authors:** Justin N. Vaughn, Walid Korani, Joshua C. Stein, Jeremy D. Edwards, Daniel G. Peterson, Sheron A. Simpson, Ramey C. Youngblood, Jane Grimwood, Doreen H. Ware, Anna M. McClung, Brian E. Scheffler

## Abstract

The genetic basis of general plant vigor is of major interest to food producers, yet the trait is recalcitrant to genetic mapping because of the number of loci involved, their small effects, and linkage. Observations of heterosis in many crops suggests that recessive, malfunctioning versions of genes are a major cause of poor performance, yet we have little information on the mutational spectrum underlying these disruptions. To address this question, we generated a long-read assembly of a tropical *japonica* rice (*Oryza sativa*) variety, Carolina Gold, which allowed us to identify structural mutations (>50 bp) and orient them with respect to their ancestral state using the outgroup, *Oryza glaberrima*. Supporting prior work, we find substantial genome expansion is the *sativa* branch. While transposable elements (TEs) account for the largest share of size variation, the majority of events are not directly TE-mediated. Tandem duplications are the most common source of insertions and are highly enriched among 50-200bp mutations. To explore the relative impact of various mutational classes on crop fitness, we then track these structural events over the last century of US rice improvement using 101 resequenced varieties. Within this material, a pattern of temporary hybridization between medium and long-grain varieties was followed by recent divergence. During this long-term selection, structural mutations that impact gene exons have been removed at a greater rate than intronic indels and single-nucleotide mutations. These results support the use of *ab initio* estimates of mutational burden, based on structural data, as an orthogonal predictor in genomic selection.

**Significance Statement:** Some crop varieties have superior performance across years and environments. In hybrids, harmful mutations in one parent are masked by the ancestral alleles in the other parent, resulting in increased vigor. Unfortunately, these mutations are very difficult to identify precisely because, individually, they only have a small effect. In this study, we use long-read sequencing to characterize the entire mutational spectrum between two rice varieties. We then track these mutations through the last century of rice breeding. We show that large structural mutations in exons are selected against at a greater rate than any other mutational class. These findings illuminate the nature of deleterious alleles and will guide attempts to predict variety vigor based solely on genomic information.

## Introduction

Though the details vary substantially, most major crops have undergone a dramatic reduction in genetic diversity relative to their wild progenitors due to domestication and breeding [1]. In the case of rice, this reduction is compounded by self-pollination. These early events have had a substantial impact on the contemporary population’s mutational load or its “cost of domestication”. Strong selection on a few key domestication loci and long-term reductions in population size resulted in the expansion and even fixation of many mildly deleterious alleles [2]. Rice was one of the earliest models used to examine and demonstrate this cost at the sequence level [3].

A central task of modern breeding is purge to these deleterious alleles [4]. Historically, this purging has been accomplished indirectly through extensive crossing and phenotypic selection. Genomic data offers a complementary (or perhaps wholly alternative) way of explicitly identifying the alleles most likely of be deleterious [5]. In effect, with the knowledge of all deleterious mutations in hand, breeders could dramatically accelerate genetic gain through targeted crosses and much larger populations afforded by marker-based breeding value assessment of progeny [4].

While the theoretical basis for the cost of domestication is strong, few studies have investigated how this cost is manifested in long-term, realistic agronomic settings. In addition, large structural mutations (SVs) are likely to have the most dramatic phenotypic effects [6]. Indeed, they account for a disproportionate number of discovered causal variants [1,4]. Yet these events have been recalcitrant to the short-read sequencing deluge. Recent work in rice using 3,000 re-sequenced rice genomes shows a clear depletion in coding-sequence indels, a hallmark of inbreeding and selection [7]. Though this work used an extensive SV-calling pipeline, false-positive and false-negative rates were still in excess of 10% and often much greater for tandem duplications and inversions [7,8]. More importantly, it remains unclear if the segregating structural variants are present because they are neutral or because deleterious alleles have not yet been fully purged.

The US rice industry traces its commercial production along the southeast coast to the 1700s. Since *Oryza* spp is not indigenous to the USA, rice varieties from around the world were imported and evaluated for production potential through the 1920s. Currently, approximately 80% of the 1 M ha of rice grown in the USA are planted in the Mid-South region along the Mississippi River and Gulf Coast and utilize tropical *japonica* germplasm (https://www.nass.usda.gov/). Early breeding efforts in the USA focused a relatively narrow genepool of tropical *japonica* germplasm that possessed the combination of agronomic and grain quality traits desired by the domestic industry. From this US-bred material, we generated a long-read assembly of a foundational tropical *japonica* variety, Carolina Gold Select (Pi 636345) (shortened to “CarGold”, in the following text and figures). This assembly was used to generate a high-confidence set of SVs spanning the genome. In addition, we resequenced 166 varieties representing USA rice breeding efforts over the last century. Of these, we focused on 101 varieties that have been selected for in a comparable environment and have had documented gene flow based on robust pedigree information. Together this subset allowed us to characterize the full mutational spectrum across a well-defined time course of breeding and selection in the USA.

## Results and Discussion

### O. sativa genome size continued to increase after the temperate/tropical split due to a small set of retrotransposons

Large structural mutations have resulted in genome expansion in the *Oryza sativa* lineage (9). TE activity – a major driver – appears to have diminished although there is evidence for continued transpositional activity at low frequency in contemporary breeding material (10). Given the repetitive nature of TEs, their activity can be very difficult to assess using short-read sequencing technology. Long-reads, which can span the critical ~12 kb threshold of full-length retrotransposons, have proven central to characterizing TE events at sequence-level resolution. In order to identify TE events and additional complex SVs, we employed PacBio long-read technology to generate a *de novo* assembly of a foundational US rice variety, Carolina Gold (“CarGold”). In addition to its significance to US rice breeding, CarGold is representative of the major tropical branch of the *O. sativa japonica* subpopulation.

The CarGold long-read assembly resulted in 208 contigs with an N50 of 12.88 Mb (***Supplemental Table 1***). These were scaffolded into pseudomolecules using the Nipponbare temperate *japonica* reference (IRGSP-1.0.59), 97.4% of which was covered by the CarGold assembly (***Supplemental Figure 4***). In addition to high coverage, the CarGold assembly shows excellent contiguity with Nipponbare, even for centromeric content. The centromere of chromosome 6 is one exception: it appears to contain three major inversions, although we did not pursue confirmation of these given that the internal content appears to be contiguous and they represent gene poor regions outside the scope of the study.

To characterize >50bp SVs, CarGold, Nipponbare, and *O. glaberrima* chromosomes were aligned. Using the *O. glaberrima* outgroup, the ancestral state of an indel between *japonica* subtypes could be inferred (***Figure 1***A and (9, 11)). As expected, the majority of events were in the outgroup relative to *O. sativa*. Because they cannot be inferred, such outgroup events were removed in the following analyses. All inferred events were compared with SVs identified using CarGold PacBio reads directly aligned to Nipponbare reference and called with pbsv (https://github.com/PacificBiosciences/pbsv, v2.2), currently the best SV caller for long-read data (12). We expected far fewer events to be present in our set since we require stringent border alignments and outgroup alignment. Indeed, whereas pbsv calls 18,238 SVs, we called 5,571 SVs (***Supplemental File 4***). 82% of the SVs we inferred overlapped pbsv SVs. Of those that did not, 92% of involved insertion of 5% or more of the total SV. Manual curation revealed that the insertion was generally present in aligned reads but either the SV was in a hemizygous state or a deletion occurring in conjunction with the insertion (see below) had apparently disrupted pbsv’s ability to call the SV.

**Figure 1:**
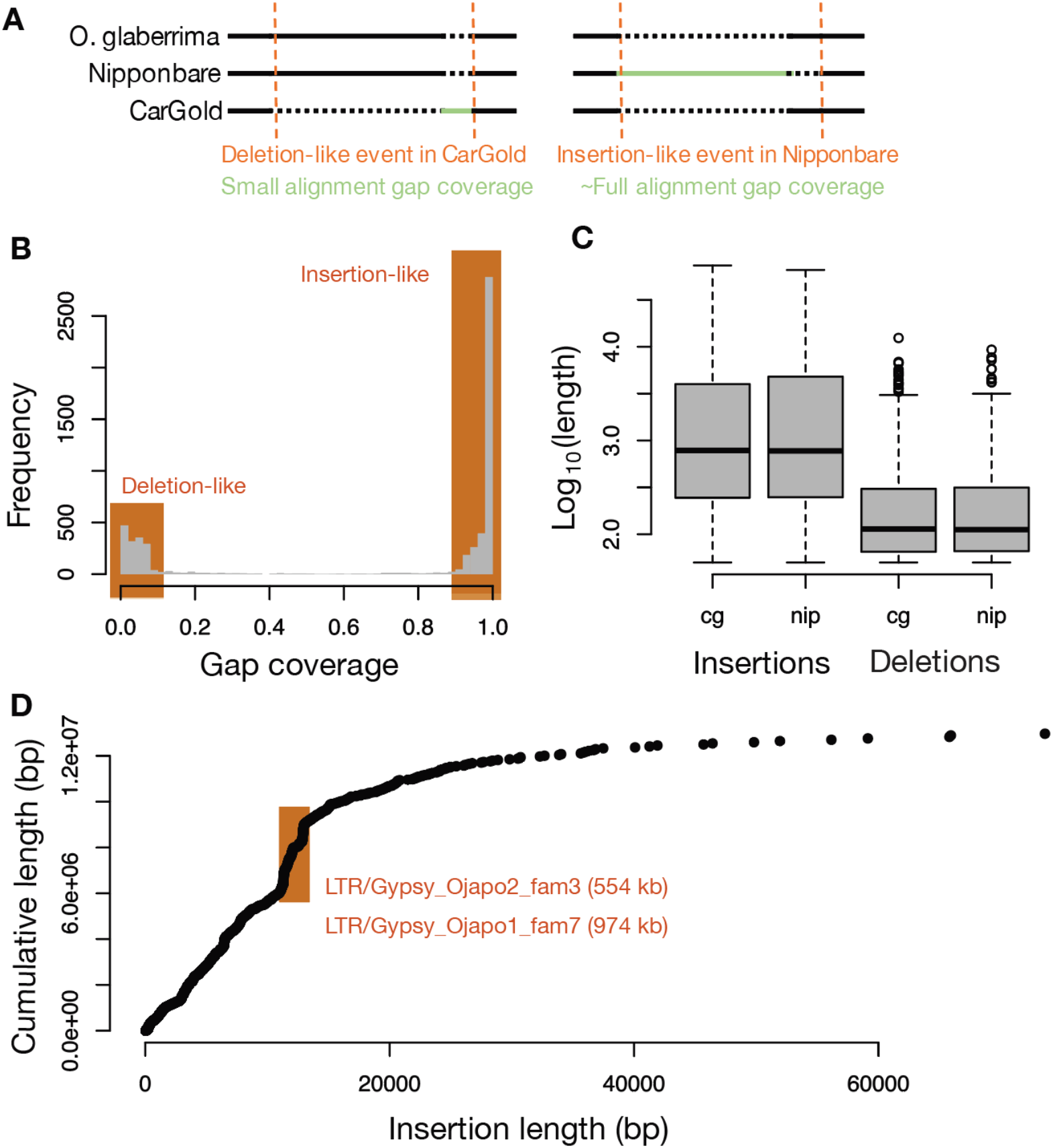
Insertions and deletions between temperate and tropical japonica references. A) Schematic illustrating the insertion/deletion orientation method for indel characterization. Any event found only in *O. glaberrima* is ambiguous and ignored. B) Distribution of gap coverage values across all events analyzed. As indicated, insertions and deletions are defined as events with a gap coverage of >95% or <5%, respectively. C) Boxplot depicting the log-transformed size distribution of events broken out by type and variety. D) Scatterplot showing each insertion event length and cumulative length after addition of that event to all shorter events. TEs contributing to rapid changes in total inserted sequence are indicated by orange window.

Most events we inferred were, as expected for a double-strand break repair process (13), a mixture of inserted and deleted bases, although generally one of the two appear to dominate an event and insertion-like events are much more frequent (***Figure 1***B). In addition to being more frequent, the insertions involve more bases on average (***Figure 1***C). The length profiles are consistent across the two varieties (***Figure 1***C). Summing across all events indicates a net gain of ~6 Mb in both lineages. As our inference methodology was conservative, this value represents a lower bound on the net gain.

The contribution of particular length-classes was not uniform (***Figure 1***D). While we observed some very large events (>40 kb), these have very little impact on the cumulative length of inserted sequence (~13 Mb). The most impactful length class was centered around 11 kb, wherein two well-defined “step-changes” were observed (***Figure 1***D). Using a full-length TE database generated specifically for this study (see Methods), we annotated the TEs within all insertions and observed a clear enrichment of two Gypsy retrotransposon families within the step-changes (***Figure 1***D). Together these two families account for ~1.5 Mb (or 11%) of inserted sequence (***Supplemental File 3***). An additional set of Copia elements, ~ 6k bp in length, accounts for another 835kb (or 6%).

### Most insertions are tandem duplications likely generated via patch-repair

While TEs, as a whole, account for 28% of the total length of added DNA, they only directly account for 9% of the total events. Though limited by short-read methodologies, prior work in rice identified that tandem duplications accounted for at least half of all new insertions >10 bp (11). These events did not appear to result from replication slippage, but DNA repair of adjacent nick sites as indicated in (11, 14). The CarGold assembly allowed us to extend prior results, which were limited to primarily <100bp events, to those SVs described above that are 10 to 100-fold longer.

We calculated a *d* metric for CarGold vs Nipponbare indels (**Figure 2**), which is a numerical description of the ambiguity in alignment between ancestral and derived states (15). Direct integration of ectopic DNA has a *d*=0. Events such as replication slippage, which should depend on small annealing sites, have a *d* greater than the length of the indel. In perfect tandem duplication, *d* equals the exact length of the indel. Our results are complementary to prior observations (11). Importantly, deletions do not conform to the 1:1 relationship seen for insertions, indicating that tandem duplication is much more common than tandem removal, the later likely occurring through unequal crossing over. This asymmetry also supports our ability to accurately infer indel ancestral state using *O. glaberrima* as an outgroup.

**Figure 2:**
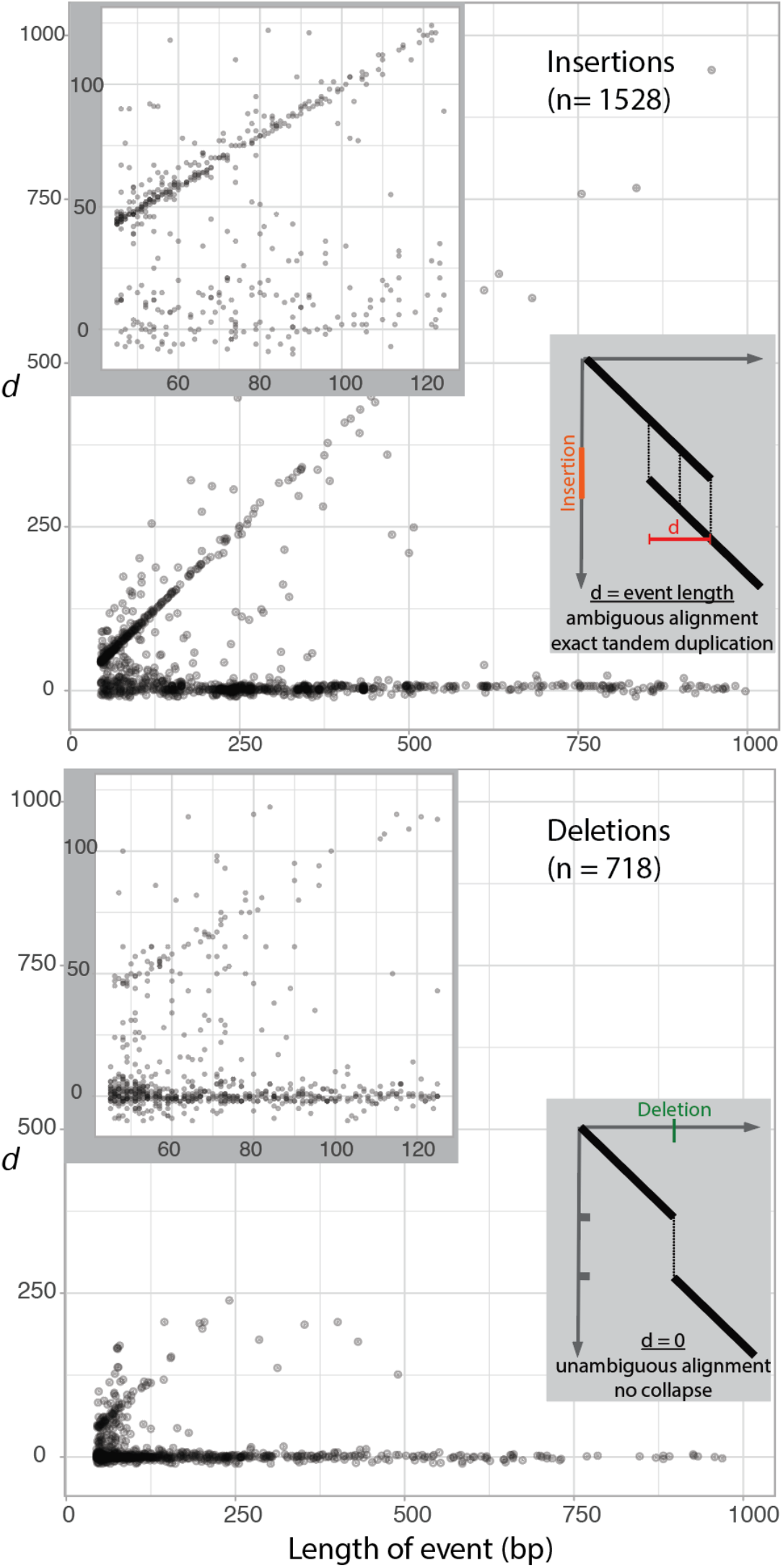
Tandem bias in insertions versus deletions between Nipponbare and CarGold. Indel size is plotted against the *d* metric for each insertion and deletion satisfying our alignment and inference criteria. Upper corner insets show only indels <125bp. Lower corner insets illustrate how *d* is calculated; alignments between ancestral and derived sequence are depicted as simplified dotplots.

Because we examined much longer mutations, we were able to identify an upper limit to the tandem duplication mechanism. As indicated in **Figure 2**, the relative proportion of tandem duplications declines as events exceed 125 bp. This proportion starts at roughly 70% of events and falls to 5% by 200 bp, based on a sliding window analysis per 40 insertions (not shown). Interestingly, the proportion of tandem duplications appears to remain at 5% out to the longest values used in the analysis (15 kb). Indeed, we observed perfect tandem duplications in excess of 1 kb, with the longest being 4.9 kb.

Across both varieties, we examined 5,571 >50bp mutations. As described in the Introduction, the presence of many of these events are likely to be a consequence of the domestication bottleneck. Given increased population size and improved measurement accuracy of yield under modern breeding, a signature of selection pressure to purge these possibly deleterious alleles should be detectable. To interrogate this relationship in an agricultural context, we turned to the US rice germplasm collection, which has representative sampling of released varieties spanning the last century of rice breeding in the US.

### Admixed introductions are followed by targeted breeding efforts for long and medium grain markets

One-hundred and sixty-six rice varieties developed or used in US rice breeding were sequenced as Illumina 100-150 bp, paired-end reads (***Supplemental File 1***). For population analysis, single nucleotide variations (SNPs) were called relative to the Nipponbare reference. This set of ~11.6M SNPs was cross-filtered with SNP data derived from ~3K other rice varieties (16), resulting in ~3.9M intersecting SNPs.

For examining the population structure in the US sample, we further filtered full SNPs by retaining those SNPs with low pairwise LD between one another (17). Thirty randomly sampled representatives from each rice subpopulation found in pre-existing data on ~3k rice varieties (16) were used as training populations to infer the proportional origin of each US variety (**Figure 3**A).

**Figure 3:**
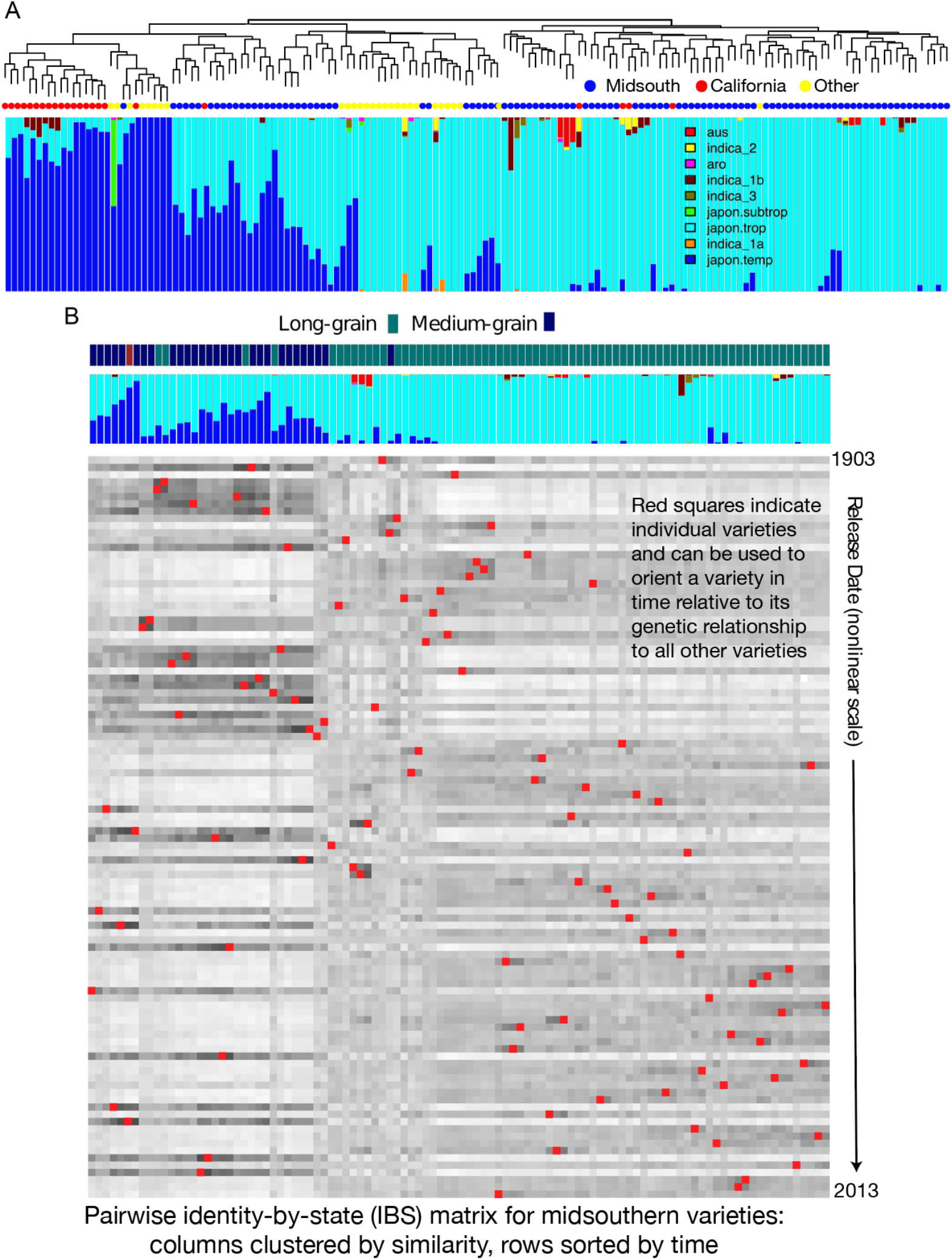
The genetic structure of US rice breeding varieties. [Shown on next page] A) The population structure of sequenced US germplasm samples in this study. The dendrogram represents hierarchical clustering based on akinship matrix derived from the low LD marker set (See Methods). Highly diverse material is shown disconnected on the top right. Bar plots, based on the same markers, indicate the proportion of genetic content of each individual that can be assigned to a set of known rice subpopulations. Colored dots represent the general location/program from which material was derived. B). Heatmap of the centered-identity-by-state (IBS) between each variety in the analysis and all other varieties, both along the column and row axis. Columns are clustered based on similarity. Rows are ordered explicitly by date of release. The red squares connect a variety’s clustering position with its release-date position. Seed type is indicated along the top, clustering axis. C. The mean and standard deviation of pairwise IBS matrix values after dividing each window of 30 varieties into the two clusters that maximize the similarity between members based on IBS profiles across the entire population. The release date axis applies to both B and C, but in C starts at the 25^th^ variety because a sliding window of 25 varieties was used in C with a +1 increment. The black line (with units on lower axis) depicts the average distance between individuals across the two clusters.

US rice varieties with substantial temperate *japonica* content are primarily derived from California breeding program. MidSouth varieties are generally tropical *japonica,* but a substantial fraction exhibit admixture between the two *japonica* types. Grain type appears to be the underlying trait that differentiates admixed MidSouth varieties from purely tropical varieties (**Figure 3**B). While >8% temperate admixture generally are associated with medium-grain varieties, there are clearly exceptions: in some cases ~50% admixture can exhibit long-grain type, whereas <2% can still have medium-grain type. Though varieties tend to be genetically distinct by grain type, admixtures are clearly present (**Figure 3**B), indicating gene flow between the subpopulations as a result of breeding efforts.

We examined population structure of varieties over the course of the last century by ordering each Midsouthern variety based on year of release (**Figure 3**B). Red dots indicate a variety’s release date relative to its genetic relatedness to all other varieties. Generally, varieties are more closely related to varieties released within the same time period, as expected for breeding populations. Initially, admix populations derived from foreign sources were introduced in the early 1900s from within which selections were made that were adapted to the southern USA. The majority of varieties that exhibit unusual seed type given their population assignments appear during this time period. Controlled crosses were implemented in 1929 with a focus on long grain development for the next 25 years. With this market established, there was an emphasis to also develop a medium grain market with several released in the 1950s, However, after 1960 over 70% of the new releases were long grains. Thus, over time, the total population has diverged through breeding and selection and resultant gene pools narrowed targeted to long and medium grain market classes.

### Structural mutations are being purged from the germplasm

This historical collection of varieties allowed us to further examine the impact of different mutational classes in a long-term breeding context. Indeed, selection against SVs is supported by our initial observation of one SV per ~424kb of exonic space versus ~222kb of intronic space (described below). Though biased TE distribution may be a factor, TE-mediated events are the minority and this ~2-fold bias likely reflects the historical purging of SVs in exons. Comparable results were recently observed in *O. sativa* using short-read SV inference (7). Yet it remains unknown if the SVs that we do observe escaped negative selection because they are neutral or if selection has been constrained by linkage and narrow genetic bottlenecks related to domestication (see Introduction). To examine this question, we explored changes in assorted mutational classes.

Using release-date as a surrogate for agronomic fitness, we estimated the effects of the large structural variants discovered above as well as a random subset of SNPs, for which we were able to define an ancestral state using *O. glaberrima* as above. Alleles were called based on read contiguity across these pre-ascertained sites (see Methods). Read coverage across all varieties was not significantly correlated with reference allele calls (*p*-val = 0.08) and a linear regression model indicates that, if there was a significant relationship, it would only account for 1.7% of our variance in genome-wide genotyping (***Supplemental Figure 10**A*). We additionally used 207x CarGold short-read sequencing data to independently assess false-negative allele calls among those SVs that were segregating in Midsouthern varieties. Assuming the SVs defined above are all true, no CarGold allele should be called as the Nipponbare reference. We downsampled reads randomly to reflect the interquartile coverages across the US varieties: 22x, 31x, and 40x, single copy coverage. A false-negative rate of 10% across all events (***Supplemental Figure 10**A*) is rivaled only by the ability to call small deletions using short-read data in prior studies (7, 8). Indeed, manual curation suggests “miscalls” were generally the result of heterozygous-appearing loci (see above), potentially resulting from tandem duplication. Since our approach is referenced biased, these SVs will inevitably get called as Nipponbare alleles. Thus, our effective false-negative rate is likely to be substantially lower than 10%.

All allele calls (SNPs and SVs) relative to the ancestral state were linearly regressed on release-date to give the rate of change across all samples (18). This methodology is comparable to that used previously (19), although, because we are comparing groups of variants spread relatively evenly through the genome (***Supplemental Figure 6***), population structure should not be a confounding factor. SNPs that produce synonymous amino-acid substitutions, relative to their ancestral state, using both Nipponbare and CarGold-derived variants, had an average effect of essentially zero (not shown). Unfortunately, it is difficult to assess variants that have a major impact on gene structure from an ancestral perspective because the reference, Nipponbare, was used to determine that gene structure. For example, ancestral deletions that remove a gene in Nipponbare will be missed because the resultant affected region simply lacks the gene to be annotated (20). To that end, we focused only on mutations in which the Nipponbare reference matches the ancestral state and so by inference originated in the CarGold lineage. Within this set, we focused on six major mutational classes: exonic, intronic, and intergenic SVs and SNPs that introduced stop-codons, non-synonymous amino acids, or have no effect on protein coding (synonymous) (**Figure 4**).

**Figure 4:**
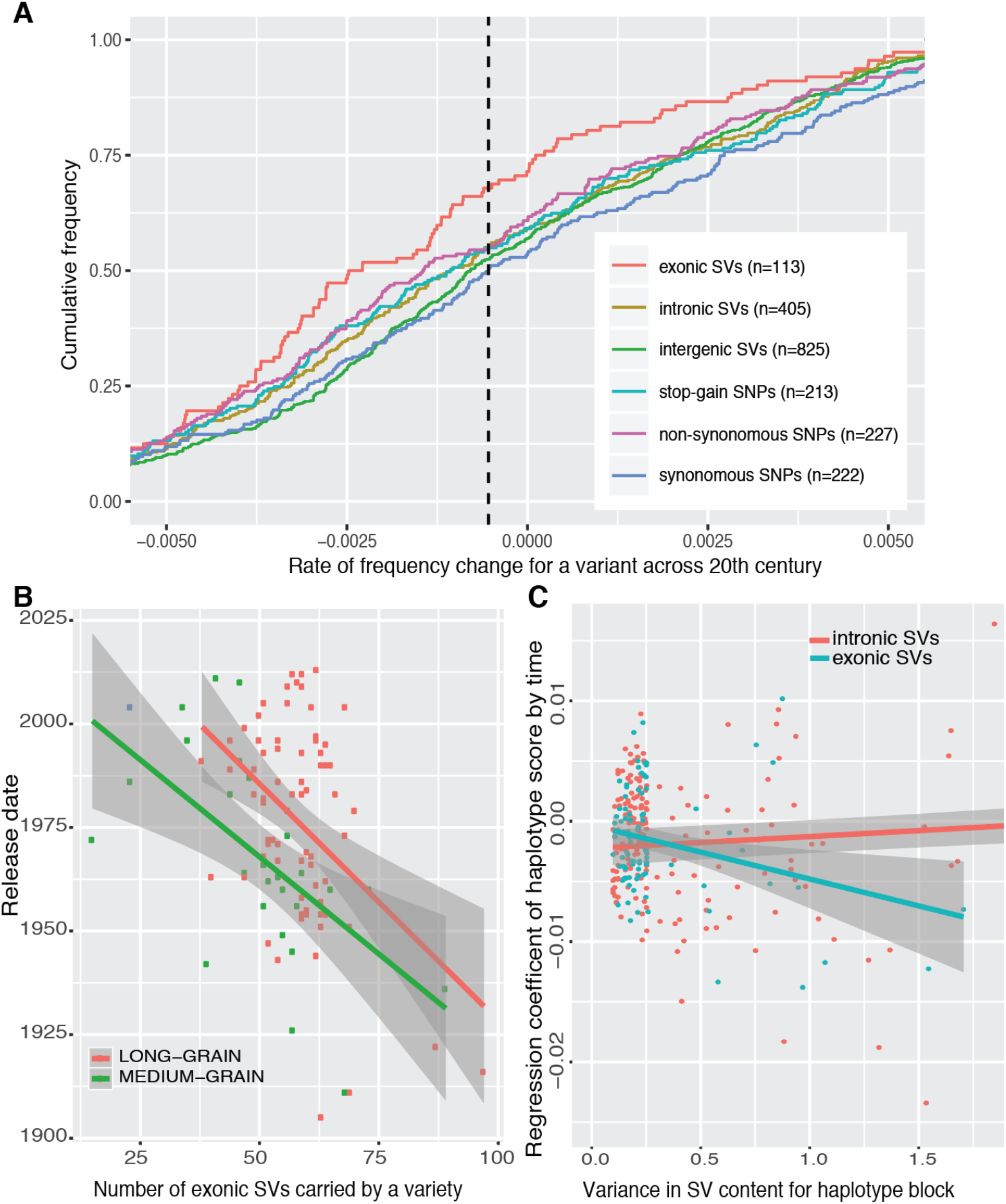
The rate of change in functional variant classes across a century of plant breeding. Rate estimates are reported only for mutations that have occurred in CarGold, not Nipponbare (see Text). A) For each variant class, the rate is plotted against that class’s cumulative frequency. The farther a class is shifted to the left of 0, the more rapidly it has declined on average. B) Plot and linear fit of release date versus the total count of allele’s that disrupt an exon in each variety for each grain type. C) Relationship between change in haplotypes scored by number of exonic indels and the variance in this score (see Figure 5). The y-axis represents, for each LD block, the regression coefficient of a linear model between haplotype score and the release date of the variety containing that haplotype. The x-axis represents the variance for those haplotype scores. In B and C, the shading around the regression line represents the 95% confidence intervals in combined intercept and slope estimates.

All median rates for each class are shifted to the left of synonymous SNPs. Given that these are CarGold-derived variants, the negative shift for synonymous SNPs, indicated by dotted line in **Figure 4**A, likely reflects a general selection against unadapted CarGold haplotypes. Though the signal is small in some cases, the median values for each class are roughly ordered based on the expected impact on open reading frame disruption (**Figure 4**A). Least significant difference test (implemented in “agricolae” library in R with p.adj = “fdr”) supports the following grouping of effect classes: Synonymous SNPs and intergenic SVs shared similar profile, particularly with regard to the most negative rates. Intronic SVs and both non-synonymous and stop-gain SNPs are effectively equivalent with median values shifted to the left of low effect mutations. Exonic SVs are substantially more negatively shifted than any other group. Within this group, we did not observe any bias with regard to length or deletion/insertion status (***Supplemental Figure 5***), supporting the expectation that any event >50 bp has an equivalent impact on gene disruption. Events had comparable density across chromosomes (***Supplemental Figure 6***), and results were essentially unaffected when chromosomes with the most divergent patterns (Chr 6 or 11) were removed from the analysis (not shown). If intergenic read mapping is less accurate than exonic mapping, a negative relationship between a variety’s year-of-release and its genetic distance to the reference could skew results: allele calls would be falsely biased toward the derive mutation in intergenic regions of newer varieties. In fact, we observe the opposite: a marginal positive relationship between year-of-release and distance to reference (***Supplemental Figure 9***), indicating that the signal of selection against gene disrupting SVs is, if anything, slightly diluted. There was also no discernable relationship between a mutation type, its rate of change, and its false-negative genotyping rate described above (***Supplemental Figure 10**B*). Lastly, in the case of true selection, rates based on linear regression will be strongest for variants that have a minor allele frequency of 0.5 across the entire sample. Derived allele frequencies were different between mutational classes, as expected for alleles that have a different impact in general (***Supplemental Figure 11***). Exonic SVs exhibit relative enrichment between 0.15 and 0.3. Again, this indicates that exonic rate estimate is biased toward weaker (not stronger) values.

Based on these results, we expected to be able to partially predict a variety’s year-of-release from the sum of exonic SVs that it contained. We observed that the relationship between reduced exonic SVs and later release date holds for both grain-types, in spite of the fact that these events are all CarGold-derived due to reasons discussed above (**Figure 4**B). In the long-grain material, the relationship is heavily driven by the exonic SV content of the oldest varieties and would likely be only marginally useful in contemporary selection. That said, these events represent only a fraction (20-30% by our rough estimate) of the exonic SVs segregating at >10% in this population. Given the strength of signal we observe (R^2^=0.24, *p*-val =1.63e-06), we predict that the addition of those data would substantially improve prediction accuracy (see Conclusions). Moreover, these predictions do not include the impact of other mutational classes as well.

Estimates of the effects of any variant will be complicated by linkage: a locus containing a beneficial and deleterious allele in positive phase are effectively invisible to selection, assuming these alleles are of roughly equivalent effect. This relationship between linkage and selection should be apparent from our data as well. To test this (and to facilitate locus-specific selection analysis below), we defined LD blocks for the combined MidSouth population using a downsampled set of ~60k SNPs (**Figure 5**). (Increased coverage had only a minor impact on refining LD blocks further, see Methods.) We counted the number of derived exonic structural mutations present in each haplotype within a given LD block. We estimated a regression coefficient between this score of each haplotype and the year-of-release of the variety in which it was present. Our hypothesis was, for exonic SVs, that the LD blocks with the largest variance in haplotype score would exhibit the most negative regression coefficients. Intronic SVs were used as a conservative “neutral” control, given that they are positionally correlated and show some bias (**Figure 4**A). Exonic SVs do exhibit this relationship relative to intronic SVs (**Figure 4**C). Due to ascertainment and probably selection, the majority of our data contains only 1 exonic SV per LD block and thus we have few exonic data-points with substantial variance. Still, the regression coefficient for exonic indels is statistically significant (p-val < 0.01) as indicated by 95% confidence intervals (**Figure 4**C), supporting the idea that haplotype context has an impact on an individual allele’s ultimate trajectory (18, 21).

**Figure 5:**
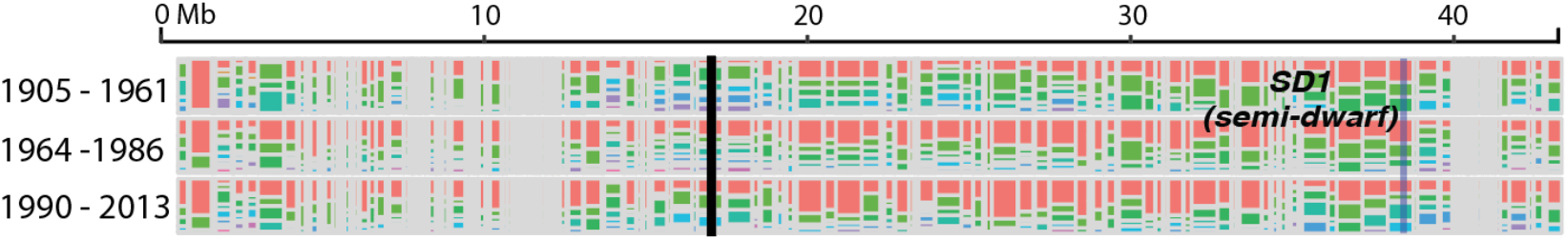
The select-ome of US rice breeding. Chromosome 1 is shown as representative. The chromosome is divided into LD blocks and haplotypes within those blocks are colored based on red being the most frequent across the three time periods defined on far left. Semi-dwarf locus, a known target of selection, is labeled. Sparse regions represent very small LD blocks. All chromosomes are plotted in **Supplemental Figure *8***. See github.com/USDA-ARS-GBRU/HaploStrata for fully interactive plots.

### 3% of LD blocks exhibit sweep-like changes in frequency

The difference in rates of allele frequency change between functional classes of mutations indicates that genetic drift in this rice population is not substantial enough to mask the signature of selection of certain loci. While weakly deleterious alleles appear to have a cumulative effect across the genome, highly adaptive mutations can occur against this backdrop and sweep through a population. We can track haplotypes through time (see above) and identify those with aberrant changes expected from such a sweep. Still, the identification of selective sweeps in unreplicated populations is notoriously problematic (22). Positive controls can be useful in establishing a realistic threshold for selection (23, 24). Starting in the late 1960s, the semi-dwarf phenotype was introduced into Asian and then US breeding programs, often increasing yields in modern agronomic environments by 50% or more (25). Though multiple alleles exist, the major source in our population is the Taichung Native 1 allele derived from an *indica* source (26).

For all haplotype blocks in our sample, we partitioned varieties into 3 time periods: early (1905-1961), middle (1964-1986), and late (1990-2013). The *sd1* block indicates a rapid increase in two nearly identical haplogroups by 23% from middle to late eras (**Figure 5**). Using this value as a threshold, 3% of LD blocks exhibit an equivalent or greater change in single haplotypes from one era to another. We also overlaid known agronomic genes (n=46, ***Supplemental File 6***) and examined the overlap with frequency change. Though numerous genes overlapped, including the pi-ta locus that determines resistance to a number of pathotypes of the rice blast fungus (*Magnaporthe oryzae)* (27), this proportion did not deviate substantially from random expectation. Thus, many putatively selected regions have no obvious target of selection (***Supplemental File 5***). To facilitate further exploration of these regions, an interactive tool complementary to **Figure 5** is available at https://github.com/USDA-ARS-GBRU/HaploStrata

The LD block sizes in our population represent the diversity in the population, either generated by recombination or by introduction of novel haplotypes. Strong selection on rare or *de novo* mutations can produce LD blocks in later generations that are much larger than those of earlier generations (28). Such a pattern would not be immediately evident in **Figure 5**. Barring heightened selection for recombination, the rate of change in alleles and the ratio of LD block lengths in early versus late generations should be highly correlated.

By evaluating pairwise identity across early and late generations (or “HScan Index”, see Methods), we did note a general shift toward longer linkage blocks (***Supplemental Figure 2*** and ***Supplemental Figure 7***), likely reflecting the population divergence observed above (**Figure 3**) and mild depletions in diversity. The distribution also has a prominent tail, likely reflecting selected outlier loci. The differences in block lengths are associated with selected loci, as defined at the Δ23% level above, exhibiting 11% longer LD in the late versus early intervals (***Supplemental Figure 3***). Interestingly, the *sd1* locus exhibits reduction in the HScan index. This change is due to the loci originally being homogenous in early types; as *sd1* began to sweep into but not completely through the population it has actually increased diversity. Only one other locus (on chromosome 7, ***Supplemental Figure 7***) had such reduced diversity in the early era and it too shows substantial increase in diversity through time.

Soft sweeps occur when an allele appears in multiple haplotypes prior to an increase in selection efficiency (28). These events can be very difficult to detect because the change in any one of the containing haplotypes is quite small. Only one of the 367 selected LD blocks possess multiple selected haplotypes. Though this percentage is almost certainly a low estimate of soft sweeps given the sensitivity afforded by our sample size, the full scope of this research supports a model in which advantageous mutations of the same locus will likely only exist in a small number of haplotypes fit enough to rise in frequency.

## Conclusions

Current methodologies for genomic selection use phenotypic and genotypic data from a training population to predict the yield of varieties that have little or no phenotypic information (29). The accuracy of the prediction is highly correlated with the relatedness between the training and test populations, and predictions can be made from a small number of markers (30). Such findings fit well with an infinitesimal model of yield: selection models based on a single training population lack generality because an unrelated test is comprised of a very distinct sets of linked alleles. The effect of these individual alleles on yield can never be realistically estimated in a QTL-mapping context, but it may be possible to gauge their effect if evaluated as a class of mutations. Variety performance could then be predicted as a sum of these effects or as a composite model with phenotypic and genotypic data (5). In this study, we have attempted to define which mutational classes would be the most relevant to such an approach by looking at selection across a broad agronomic environment, namely, Midsouthern US varieties in the 20^th^ century. Sequence conservation scores, such as GERP, have been used previously to generate *de novo* breeding values using SNPs (5). Our results indicate that while SNPs are relevant, predictive power could be substantially improved by considering SVs, particularly those that disrupt exons. That said, we did not observe an obvious way to weigh different exonic SVs in terms of phenotypic consequences: length was not a factor (***Supplemental Figure 5***), suggesting that an additive GERP score would be inappropriate. Though not addressed here due to sample size, phylogenetic distribution of homologs may be a more useful correlate of effect size under the assumption that disruptive mutations in broadly distributed orthologs will have greater effects. Still, results using a simple 0/1 weighing scheme are encouraging (**Figure 4**B-C). Importantly, they are based solely on one reference genome annotation and SVs derived from a single, alternative assembly. Critical to the advancement of such *ab initio* prediction will be 1) the availability of enough high-quality *de novo* assemblies to capture common SVs in breeding germplasm and 2) the computational tools to annotate genes within this pan-genomic context.

## Materials and Methods

### Carolina Gold PacBio sequencing and assembly

High molecular weight DNA was extracted from young leaves using prior protocol with minor modifications (31). Essentially, young leaves, that had been flash frozen, and kept frozen at −80C, were ground to a fine powder in a frozen mortar with liquid N2 followed by very gentle extraction in CTAB buffer (that included proteinase K, PVP-40 and beta-mercaptoethanol) for 1hr at 50C. After centrifugation, the supernatant was gently extracted twice with 24:1 chloroform:iso-amyl alcohol. The upper phase was adjusted to 1/10^th^ volume with 3M KAc, gently mixed, and DNA precipitated with iso-propanol. DNA was collected by centrifugation, washed with 70% Etoh, air dried for 20 min and dissolved thoroughly in 1x TE at room temperature. Size was validated by pulsed field electrophoresis.

Libraries were prepared using PacBio SMRTbell™ Template Prep Kit 1.0, PacBio SMRTbell™ Damage Repair Kit, and prepared for sequencing using PacBio DNA/Polymerase Binding Kit P6 V2. Sequencing was performed on the PacBio RSII using PacBio DNA Sequencing Reagent 4.0 v2 and PacBio SMRT® Cell 8Pac V3. All protocols used were PacBio recommended protocols.

Raw PacBio reads (n=3,765,107; ~70x coverage) were assembled into 209 contigs, using `Canu` (v1.5) (32), and polished with `Quiver` (`smrtlink` v5.0.1 suite, now at *github.com/PacificBiosciences/GenomicConsensus*). PacBio raw reads were further polished with `pilon` v1.22 (33) using “10X Genomics” linked-reads aligned by `Longranger` v2.1.6 (*github.com/10XGenomics/longranger*). The original primary assembly consisted of 209 contigs. One contig (tig74) was determined to be a false chimeric assembly and was split into tig74a and tig74b. Repeat masking was performed as part of the `MAKER-P` pipeline (34) using custom repeat libraries, PReDa_121015_short.fasta (DNA) and TE_protein_db_121015_short_header.fasta (protein) (35, 36).

### Whole genome alignments

PacBio contigs of CarGold were assigned to Nipponbare, a temperate *japonica* variety, chromosomes (*Osativa_204_v7.0.softmasked.fa*). Exclusive pairwise relationships were determined based on the sum coverage of CarGold scaffold by collinear Nipponbare sequence. Any scaffold with a sum coverage of <65% was removed. CarGold scaffolds and their respective Nipponbare chromosomes were combined with the appropriate *Oryza glaberrima* chromosome and aligned in a chromosome-wise manner using *progressiveMauve* (37) (build-date-Feb-25-2015) with default parameters.

### Insertion/deletion inference

Indels were identified from the whole genome alignments above and were polarized relative to the outgroup, *Oryza glaberrima,* using a custom program (indelInference.pl). Columns in the whole chromosome alignments that involved >50 consecutive gaps (in any sequence) were extracted along with +/− 50 bp of flanking sequence. Gaps were analyzed further if the left and right flanking regions aligned with >90% columns being identical. If Nipponbare and CarGold shared 95% identity in the gapped region, the SV was not considered further. Alternatively, if there was variation between Nipponbare and CarGold and one matched *O. glaberrima* with >95% identity, then the event was inferred to have occurred in the non-matching sequence. This approach captured the biological reality that mutation events creating long (>50bp) SVs rarely involve only insertion or deletion of DNA but a combination of both. To that end, we also characterized the degree to which with each mutation represents a net gain or loss of DNA. The length of the entire gapped region was divided by the length of novel sequence introduced in the gap such that values approaching 0 are, in effect, deletions and values approaching 1 are insertions. (A small number of SVs with gap values between 0.49 and 0.51 were removed after manual curation indicated these “perfectly balanced” indels represent unwarranted gap openings.)

### Reference sets for TE content

An initial set of full-length TEs from the *japonica* subpopulation was retrieved from RiTE (36) (3 March 2018). This set was searched against both the Nipponbare and CarGold assemblies using *nucmer* (version 3.1) with – maxmatch. Alignments were filtered such that a full-length TE had to align to 97% of its length at a 95% identity threshold. The matching genomic region was extracted if it was not already assigned to another TE meeting the above criteria. The Nipponbare and CarGold TE sequences were combined. This combined set was searched against itself using *nucmer* with --maxmatch flag. Coordinate files were then filtered such that sequences with a reciprocal overlap of 90% were paired. *Mcl* (version 14) was used to consolidate pairs into clusters. All sequences within a cluster were aligned with mafft (version 7.307). A consensus sequence was generated using a custom program (majorityRuleConsensus.pl), such that any alignment column with A, C, G, T, or – (gap) was called based on the most frequent variant. If the gap is the most frequent, then the column is removed from the final consensus. Pipeline resulted in 7,624 TE consensus sequences (i.e. families)(***Supplemental File 2***).

### *Characterizing insertion and deletion types using* d *metric*

The *d* metric was calculated for insertions and deletions as described previously (11, 15). In brief, ancestral and derived sequences including the inserted/deleted sequence and flanking sequence (equal in length to the indel) were aligned using *BLAST* (v.2.6.0+) (38): ‘blastn - gapopen 2 -gapextend 4 -dust no’. The two major alignments were identified and the top-alignment end position was subtracted from the bottom-alignment start position, relative to the longest of the two aligned sequences (**Figure 2**)

### Short-read DNA sequencing of US germplasm sample

All sequencing libraries where generated using the Illumina TruSeq PCR-free reaction. Accession libraries were sequenced on different instruments: 24 libraries on Illumina HiSeq2500 in paired-end mode with 101 cycles, 70 libraries on Illumina HiSeqX in paired-end mode with 151 cycles, and 72 libraries on Illumina HiSeq3000 in paired-end mode with 151 cycles. Raw Illumina reads and *bbtools khist.sh* (jgi.doe.gov/data-and-tools/bbtools/) were used to assess the single-copy k-mer count for each accession, based on the main peak in the resultant kmer spectrum. Haploid genome size was also estimated as part of this analysis (***Supplemental File 1***).

### SNP-calling, merging, and pruning

Raw reads were trimmed using *trimmomatic* (39). Reads were then aligned to *Osativa_204_v7.0.softmasked.fa* using *bwa-mem,* version 0.7.17 (40). HaplotypeCaller from the GATK suite, version 4.0.8.1, was used to call SNPs and small indels (41). Each sample was used to generate a GVCF file (-T HaplotypeCaller --genotyping_mode DISCOVERY --emitRefConfidence GVCF). The combined set of GVCF files were used when genotyping the entire set under joint calling mode (-T GenotypeGVCFs). Variants were merged with those derived from a previously sequenced set of ~3k rice varieties (16); only intersecting variants were retained. 300 varieties from the 3k set, representing the 9 major subpopulations, were randomly down-sampled. This down-sampled set of 300 was combined with all US -developed varieties in this study. All singlenucleotide polymorphisms (SNPs) with a minor allele frequency less than .05 were removed. A reduced marker set was generated based on linkage disequilibria reduction as implemented in *plink* (--indep-pairwise 50 10 0.1).

### Population characterization

Population assignments and admixture estimation for US varieties was calculated using the 3k subset as the training set in *admixture’s* supervised mode (version 1.3.0,) with expected population number (K) equal to 9 (16). To analyze changes through time, all US varieties (n=166) were filtered to include only USA MidSouth sources (**Figure 3**: and ***Supplemental File 1***). Nira [PI305133] and Rexmont [GSOR 305081] were removed because of extreme divergence and poor genotyping, respectively, resulting in 101 accessions. Pruned-SNPs were further filtered from this set to remove any sites with heterozygosity >10%, as all varieties are inbred, and <20 lines without an allele call, resulting in 68,829 remaining sites. *Tassel* v5.5.50 (42) was used to create a kinship matrix from this marker set using the Centered-IBS method with maximum of 2 alleles (43).

### Calling SV genotypes

The insertions and deletions identified above were assessed for segregation across our entire resequenced population. Alignment files generated during SNP calling above were filtered to remove split and soft-clipped reads. Importantly, reads aligning to multiple locations were left in to preserve contiguity. SV-associated genomic intervals in the Nipponbare reference with >98% bases covered by one or more reads were considered support for the Nipponbare allele. If the Nipponbare interval was <20 bp (as in a CarGold insertion) then we required that at least one read span the entire interval and 5 bp flanking both sides. *bamtools/bedcov* commands used are provided (bedCovCommandForSegIndels.sh).

### Assessing allele frequency change through time

Both SV and SNP alleles were encoded as either 0 (ancestral) or 1 (derived) using information generated using the methodologies described above. The relationship between allelic state (y-values) and year-of-release (x-values) was fit to a linear model using the default *lm* function in R. Only polymorphisms with a minor allele frequency > 0.1 in Midsouthern material were used.

### Haplotype block analysis

For haplotype block assessment, the entire SNP set was filtered to MidSouth varieties (n=101 described in “Population Characterization” method) and loci with MAF >10% and heterozygote calls in more than 10 varieties were removed, resulting in 810,808 sites. In addition, GSOR 305081 (Rexmont) was removed due to poor genotyping at these loci. *Tassel* v5.5.50 (42) was used to further downsample loci by restricting distance to neighboring SNP. Starting at a distance of 20K bp, we progressively reduced distance and assessed the number of resultant haplotype blocks as a proportion total sites not in strong LD. This value plateaued at 3 kb (n = 59,575 SNPs). LD blocks were assessed using *snpldb* (v.1.2) (44), with MAF = 0.02 (at the haplotype level) and max window of 800 kbp. This 3 kb set and *h-scan* software (version 1.0: messerlab.org/resources/) was used to gauge the average length of pairwise identity tracts for each SNP position for each time period depicted in **Figure 5**.

## Data availability

Raw sequencing reads have been deposited as part of NCBI BioProject PRJNA603026. Software related to critical aspects of analysis is available at github.com/USDA-ARS-GBRU/US_rice.

## Acknowledgments

This work is supported in part by ARS projects: 6066-21310-005-00-D, 6066-21310-005-23-S, 6066-21310-005-25-S, 8062-2100-044, and 6028-21000-011-00D. USDA is an equal opportunity provider and employer.

## Supplemental Figures

**Supplemental Figure 1**: Sequencing coverage plotted against time. [SingleCopyCovAndReleaseWithMidsouth.pdf]

**Supplemental Figure 2**: Histogram of h-scan ratios [histogramHScan.pdf]

**Supplemental Figure 3**: Boxplot of h-scan ratios for putatively selected versus neutral loci [neutralVSelectedHScan.pdf]

**Supplemental Figure 4**: Dotplot of CarGold contigs relative to Nipponbare chromosomes [cgVersusNip_nucmerDot.png]

**Supplemental Figure 5**: Relationship between SV attributes and rate [lengthAndTypeVersusRate.pdf]

**Supplemental Figure 6**: Feature density of different SV classes [SVTypeDensity.pdf]

**Supplemental Figure 7**: Genomic profiles of HScan index in early versus late [LengthHScan.earlyVlate.pdf]

**Supplemental Figure 8**: Haplotype plots for all chromosomes [fullHaplotype.pdf]

**Supplemental Figure 9**: Relationship between year-of-release and genetic distance to Nipponbare reference [refBiasThroughTime.pdf]

**Supplemental Figure 10**: Genotype recall and bias assessment. A) Relationship between coverage and genome-wide genotyping as well as downsampled CarGold short-reads. B) Effect of gene location on coverage versus genotyping relationship. [CarGoldGenotypingControls.pdf]

**Supplemental Figure 11**: Cumulative frequency plots for derived SV alleles in Midsouthern sample with a frequency between 0.1 and 0.9. Each line represents the location of the SV relative to a gene. [freqDistbyGeneLoc.pdf]

## Supplemental Tables

**Supplemental Table 1**: Assembly statistics for Carolina Gold [carGoldStats.docx]

**Supplemental Table 2**: Core gene content for Carolin Gold assembly and Nipponbare reference [carGoldBusco.docx]

## Supplemental Files

**Supplemental File 1**: Master key with characteristics for all accessions sequenced as part of this study (varietyMasterKey.xlsx).

**Supplemental File 2**: TE families and consensus sequences generated in this study (consensusTEs.fasta).

**Supplemental File 3**: TE frequencies in indels (allTeInfo.xlsx).

**Supplemental File 4**: SV information, excluding sequence (allIndels.xlsx).

**Supplemental File 5**: Putatively selected loci based on sd1 threshold (selectedLoci_sd1Threshold.xlsx).

**Supplemental File 6**: Agronomically important genes (agronomicGenes.xlsx).

## References

1. B. S. Gaut, D. K. Seymour, Q. Liu, Y. Zhou, Demography and its effects on genomic variation in crop domestication. Nat. Plants 4, 512 (2018).

2. B. T. Moyers, P. L. Morrell, J. K. McKay, Genetic Costs of Domestication and Improvement. J. Hered. 109, 103–116 (2018).

3. J. Lu, et al., The accumulation of deleterious mutations in rice genomes: a hypothesis on the cost of domestication. Trends Genet. TIG 22, 126–131 (2006).

4. J. G. Wallace, E. Rodgers-Melnick, E. S. Buckler, On the Road to Breeding 4.0: Unraveling the Good, the Bad, and the Boring of Crop Quantitative Genomics. Annu. Rev. Genet. 52, 421–444 (2018).

5. J. Yang, et al., Incomplete dominance of deleterious alleles contributes substantially to trait variation and heterosis in maize. PLOS Genet. 13, e1007019 (2017).

6. D. Rodríguez-Leal, Z. H. Lemmon, J. Man, M. E. Bartlett, Z. B. Lippman, Engineering Quantitative Trait Variation for Crop Improvement by Genome Editing. Cell 171, 470–480.e8 (2017).

7. R. R. Fuentes, et al., Structural variants in 3000 rice genomes. Genome Res. (2019) https://doi.org/10.1101/gr.241240.118 (May 1, 2019).

8. M. Mahmoud, et al., Structural variant calling: the long and the short of it. Genome Biol. 20, 246 (2019).

9. J. Ma, J. L. Bennetzen, Rapid recent growth and divergence of rice nuclear genomes. Proc. Natl. Acad. Sci. U. S. A. 101, 12404–12410 (2004).

10. M.-C. Carpentier, et al., Retrotranspositional landscape of Asian rice revealed by 3000 genomes. Nat. Commun. 10, 24 (2019).

11. J. N. Vaughn, J. L. Bennetzen, Natural insertions in rice commonly form tandem duplications indicative of patch-mediated double-strand break induction and repair. Proc. Natl. Acad. Sci. 111, 6684–6689 (2014).

12. S. Kosugi, et al., Comprehensive evaluation of structural variation detection algorithms for whole genome sequencing. Genome Biol. 20, 117 (2019).

13. H. Puchta, The repair of double-strand breaks in plants: mechanisms and consequences for genome evolution. J. Exp. Bot. 56, 1–14 (2005).

14. S. Schiml, F. Fauser, H. Puchta, Repair of adjacent single-strand breaks is often accompanied by the formation of tandem sequence duplications in plant genomes. Proc. Natl. Acad. Sci., 201603823 (2016).

15. P. W. Messer, P. F. Arndt, The majority of recent short DNA insertions in the human genome are tandem duplications. Mol. Biol. Evol. 24, 1190–1197 (2007).

16. W. Wang, et al., Genomic variation in 3,010 diverse accessions of Asian cultivated rice. Nature 557, 43 (2018).

17. D. H. Alexander, K. Lange, Enhancements to the ADMIXTURE algorithm for individual ancestry estimation. BMC Bioinformatics 12, 246 (2011).

18. J. N. Vaughn, Z. Li, Genomic Signatures of North American Soybean Improvement Inform Diversity Enrichment Strategies and Clarify the Impact of Hybridization. G3 Genes Genomes Genet. 6, 2693–2705 (2016).

19. J. van Heerwaarden, M. B. Hufford, J. Ross-Ibarra, Historical genomics of North American maize. Proc. Natl. Acad. Sci. 109, 12420–12425 (2012).

20. X. Gan, et al., Multiple reference genomes and transcriptomes for Arabidopsis thaliana. Nature 477, 419–423 (2011).

21. W. G. Hill, A. Robertson, The effect of linkage on limits to artificial selection. Genet. Res. 8, 269–294 (1966).

22. T. M. Beissinger, et al., A Genome-Wide Scan for Evidence of Selection in a Maize Population Under Long-Term Artificial Selection for Ear Number. Genetics 196, 829–840 (2014).

23. J. K. Pritchard, J. K. Pickrell, G. Coop, The Genetics of Human Adaptation: Hard Sweeps, Soft Sweeps, and Polygenic Adaptation. Curr. Biol. CB 20, R208–R215 (2010).

24. T. Bersaglieri, et al., Genetic Signatures of Strong Recent Positive Selection at the Lactase Gene. Am. J. Hum. Genet. 74, 1111–1120 (2004).

25. D. S. Athwal, Semidwarf Rice and Wheat in Global Food Needs. Q. Rev. Biol. 46, 1–34 (1971).

26. B. Angira, et al., Haplotype Characterization of the sd1 Semidwarf Gene in United States Rice. Plant Genome 12 (2019).

27. K. Rybka, M. Miyamoto, I. Ando, A. Saito, S. Kawasaki, High Resolution Mapping of the Indica-Derived Rice Blast Resistance Genes II. Pi-ta2 and Pi-ta and a Consideration of Their Origin. Mol. Plant. Microbe Interact. 10, 517–524 (1997).

28. P. W. Messer, D. A. Petrov, Population genomics of rapid adaptation by soft selective sweeps. Trends Ecol. Evol. 28, 659–669 (2013).

29. A. J. Lorenz, et al., “Chapter Two - Genomic Selection in Plant Breeding: Knowledge and Prospects” in Advances in Agronomy, D. L. Sparks, Ed. (Academic Press, 2011), pp. 77–123.

30. B. B. Stewart-Brown, Q. Song, J. N. Vaughn, Z. Li, Genomic Selection for Yield and Seed Composition Traits Within an Applied Soybean Breeding Program. G3 Genes Genomes Genet. 9, 2253–2265 (2019).

31. J. Doyle, et al., A rapid DNA isolation procedure from small quantities of fresh leaf tissues (1987) (April 7, 2020).

32. S. Koren, et al., Canu: scalable and accurate long-read assembly via adaptive k-mer weighting and repeat separation. Genome Res., gr.215087.116 (2017).

33. B. J. Walker, et al., Pilon: An Integrated Tool for Comprehensive Microbial Variant Detection and Genome Assembly Improvement. PLOS ONE 9, e112963 (2014).

34. B. L. Cantarel, et al., MAKER: An easy-to-use annotation pipeline designed for emerging model organism genomes. Genome Res. 18, 188–196 (2008).

35. J. C. Stein, et al., Genomes of 13 domesticated and wild rice relatives highlight genetic conservation, turnover and innovation across the genus Oryza. Nat. Genet. 50, 285–296 (2018).

36. D. Copetti, et al., RiTE database: a resource database for genus-wide rice genomics and evolutionary biology. BMC Genomics 16, 538 (2015).

37. A. E. Darling, B. Mau, N. T. Perna, progressiveMauve: multiple genome alignment with gene gain, loss and rearrangement. PloS One 5, e11147 (2010).

38. C. Camacho, et al., BLAST+: architecture and applications. BMC Bioinformatics 10, 421 (2009).

39. A. M. Bolger, M. Lohse, B. Usadel, Trimmomatic: a flexible trimmer for Illumina sequence data. Bioinformatics 30, 2114–2120 (2014).

40. H. Li, R. Durbin, Fast and accurate short read alignment with Burrows-Wheeler transform. Bioinforma. Oxf. Engl. 25, 1754–1760 (2009).

41. M. A. DePristo, et al., A framework for variation discovery and genotyping using next-generation DNA sequencing data. Nat. Genet. 43, 491–498 (2011).

42. P. J. Bradbury, et al., TASSEL: software for association mapping of complex traits in diverse samples. Bioinformatics 23, 2633–2635 (2007).

43. J. B. Endelman, J.-L. Jannink, Shrinkage estimation of the realized relationship matrix. G3 Bethesda Md 2, 1405–1413 (2012).

44. J. He, et al., An innovative procedure of genome-wide association analysis fits studies on germplasm population and plant breeding. Theor. Appl. Genet. 130, 2327–2343 (2017).

