## Supplemental Table 2 for "Gene disruption by structural mutations drives selection in US rice breeding over the last century"

**Supplemental Table 2**: Core gene content for Carolin Gold assembly and Nipponbare reference

| **Result** | **Carolina Gold** | **IRGSP-1.0** |
| --- | --- | --- |
| Complete BUSCOs | 940 | 941 |
| Complete and single-copy BUSCOs | 912 | 917 |
| Complete and duplicated BUSCOs | 28 | 24 |
| Fragmented BUSCOs | 5 | 2 |
| Missing BUSCOs | 11 | 13 |
| Total BUSCO groups searched | 956 | 956 |
