## Supplemental Table 1 for "Gene disruption by structural mutations drives selection in US rice breeding over the last century"

**Supplemental Table 1**: Assembly statistics for Carolina Gold

| **assembly** | **count** | **sum_len** | **N50** | **min_len** | **max_len** | **med_len** |
| --- | --- | --- | --- | --- | --- | --- |
| contigs | 208 | 386,298,647 | 12,879,605 | 4,005 | 23,384,950 | 75,423 |
| unitigs | 4430 | 463,206,789 | 576,500 | 2,007 | 4,351,218 | 24,438 |
| unassembled | 198243 | 1,597,111,033 | 11,260 | 1,999 | 73,438 | 5,703 |
