## Supplementary figures and images for "Gene disruption by structural mutations drives selection in US rice breeding over the last century"

### Supplemental Figure 1

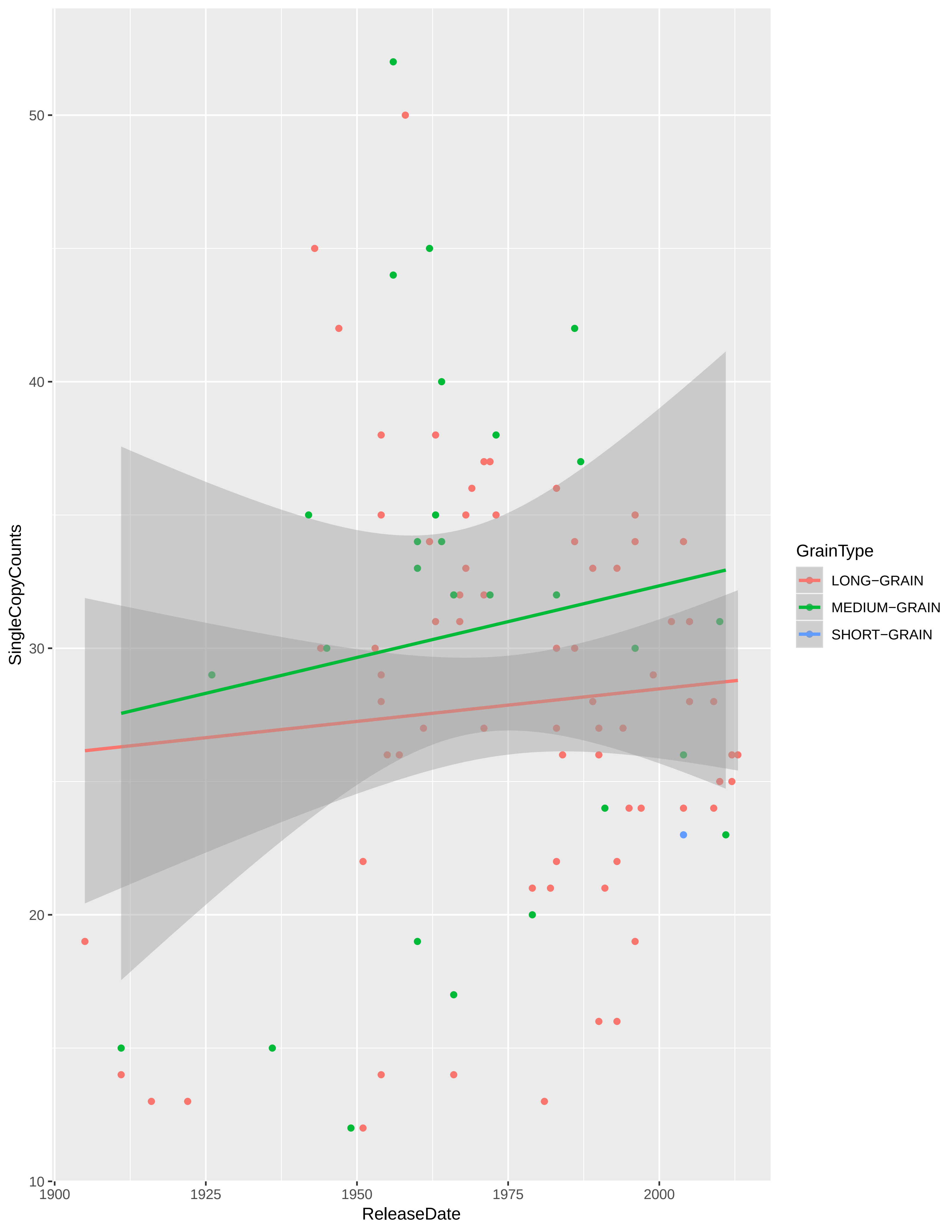

### Supplemental Figure 2

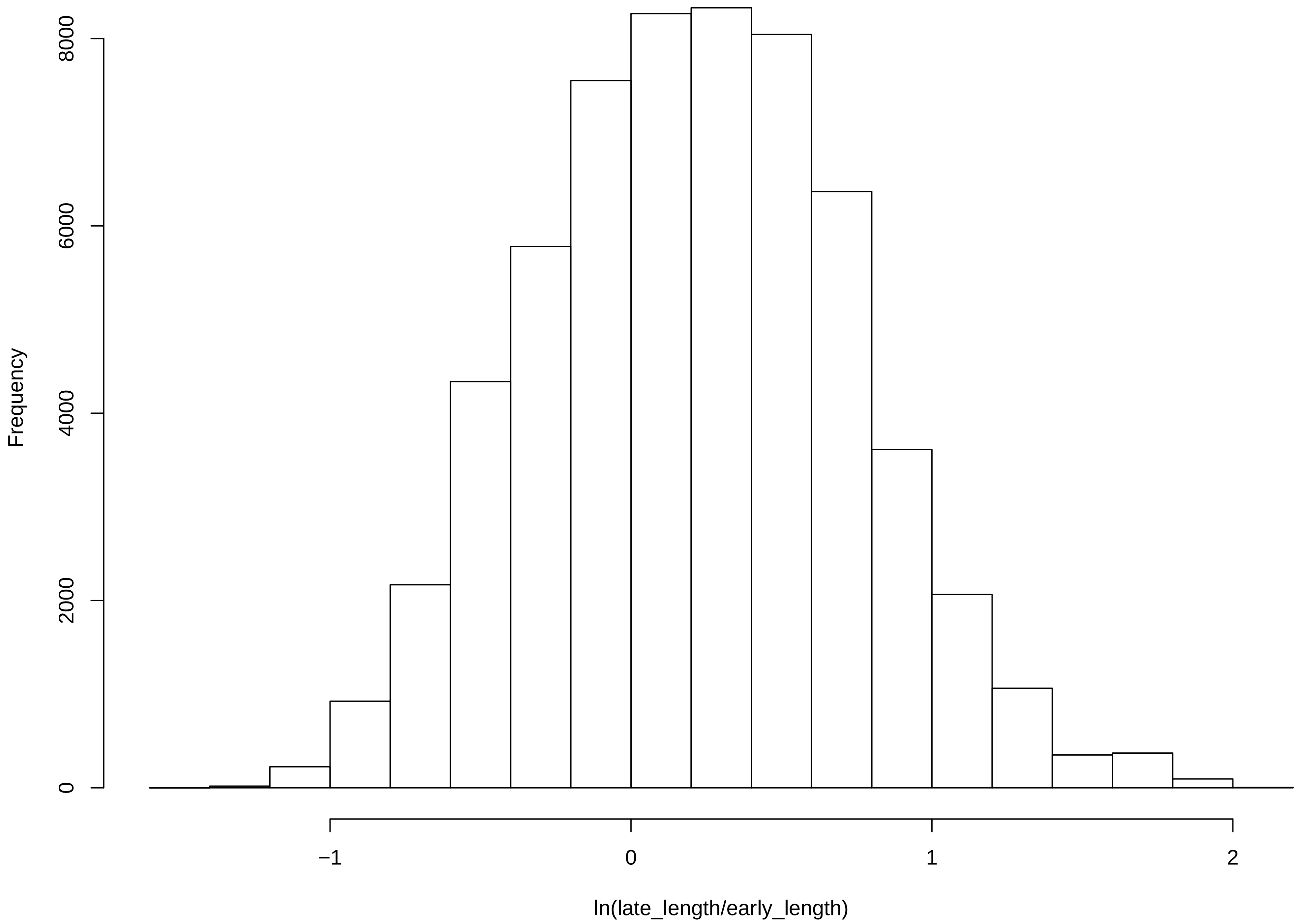

### Supplemental Figure 5

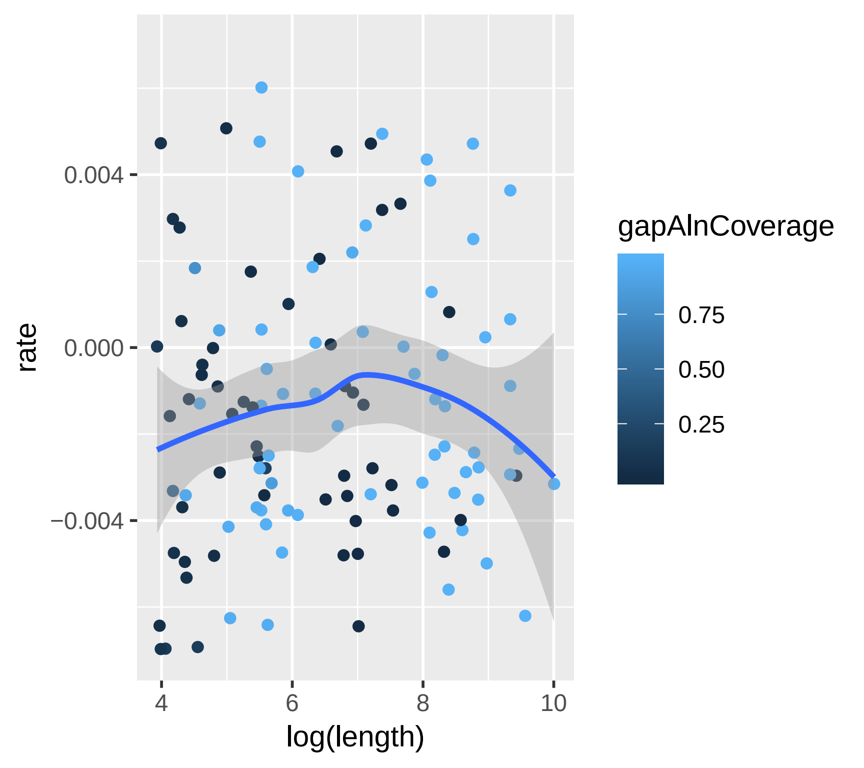

### Supplemental Figure 6

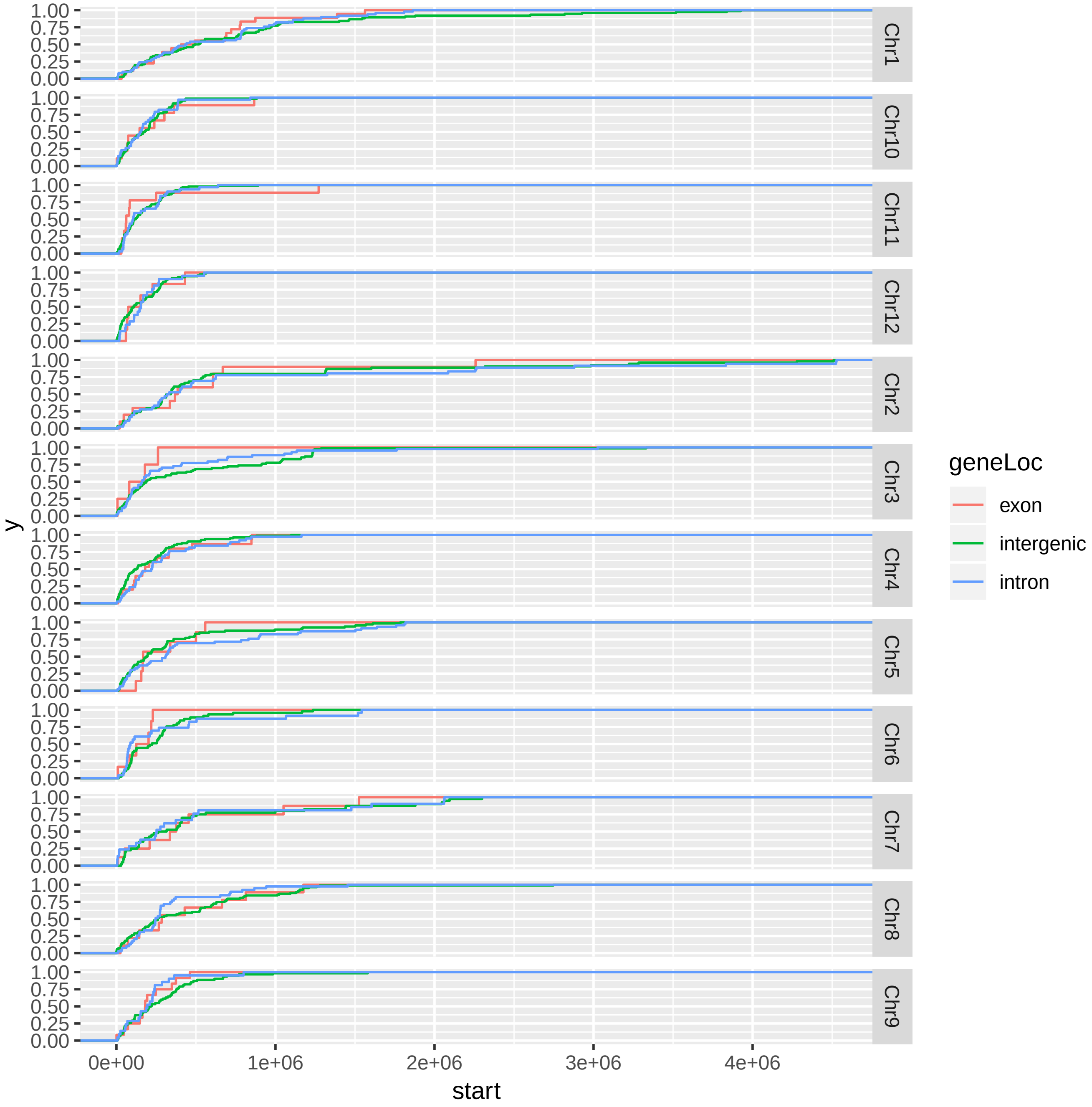

### Supplemental Figure 7

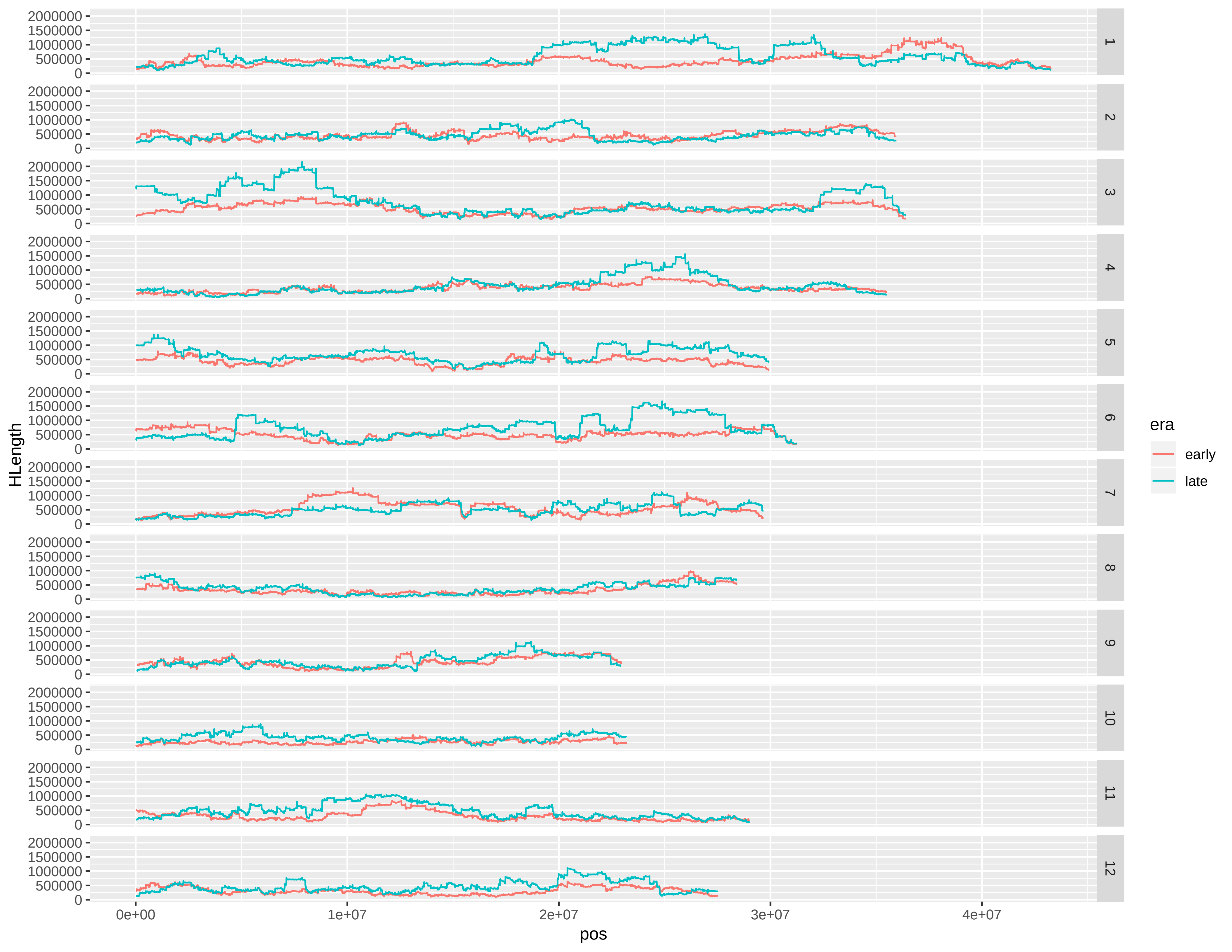

### Supplemental Figure 8

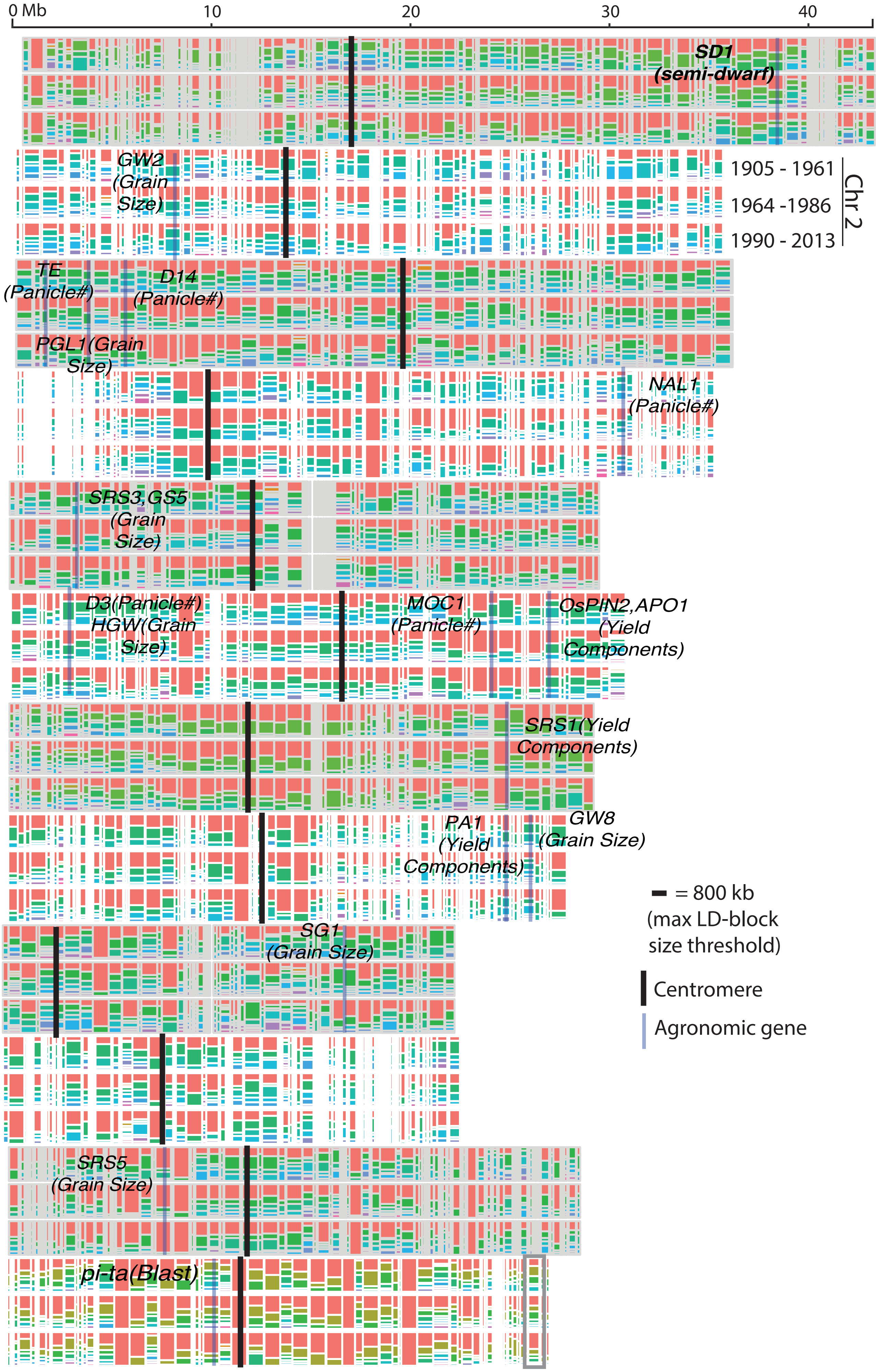

### Supplemental Figure 9

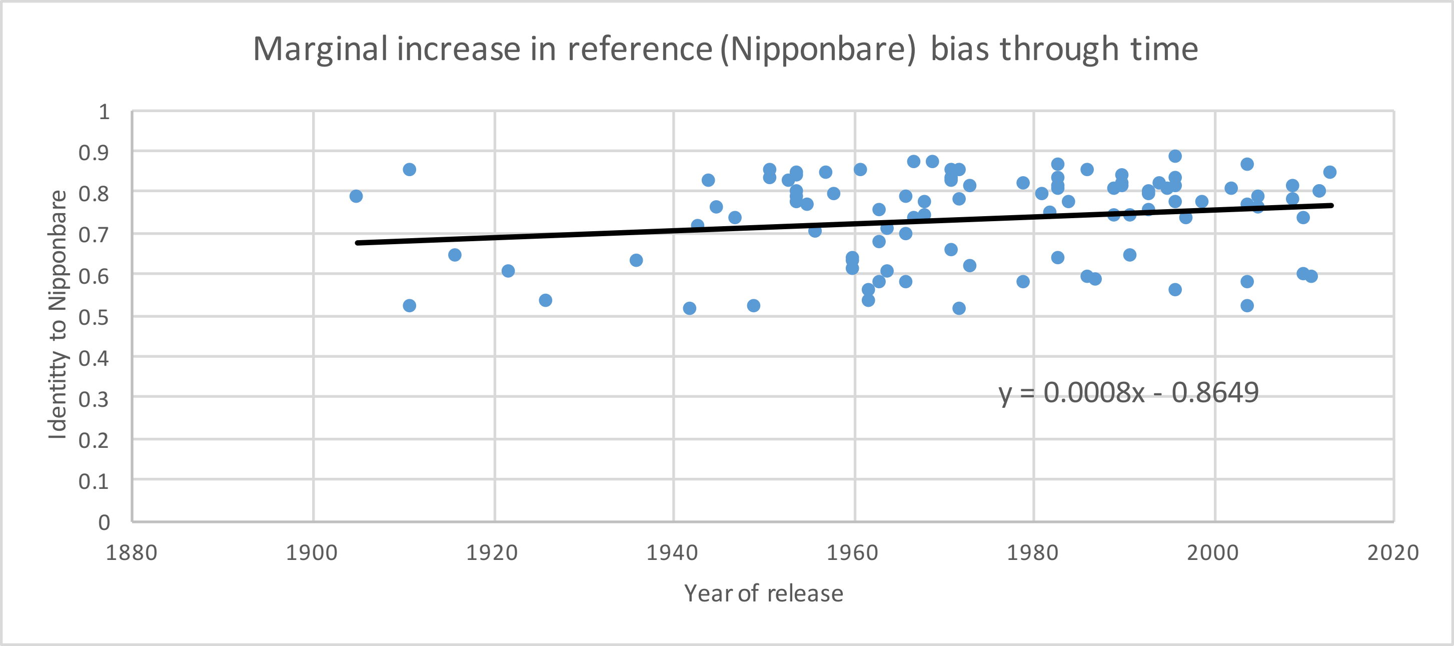

### Supplemental Figure 10

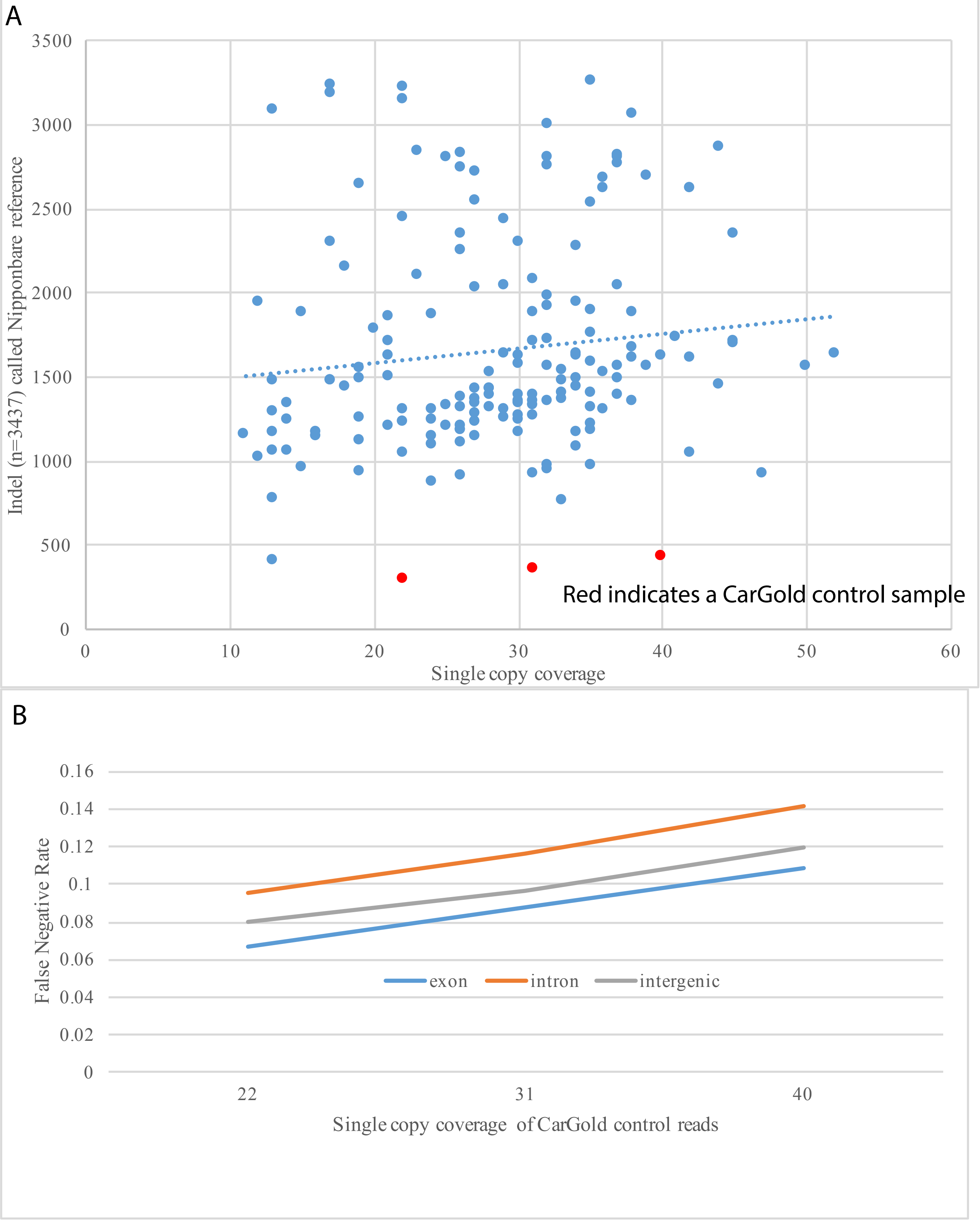

### Supplemental Figure 11

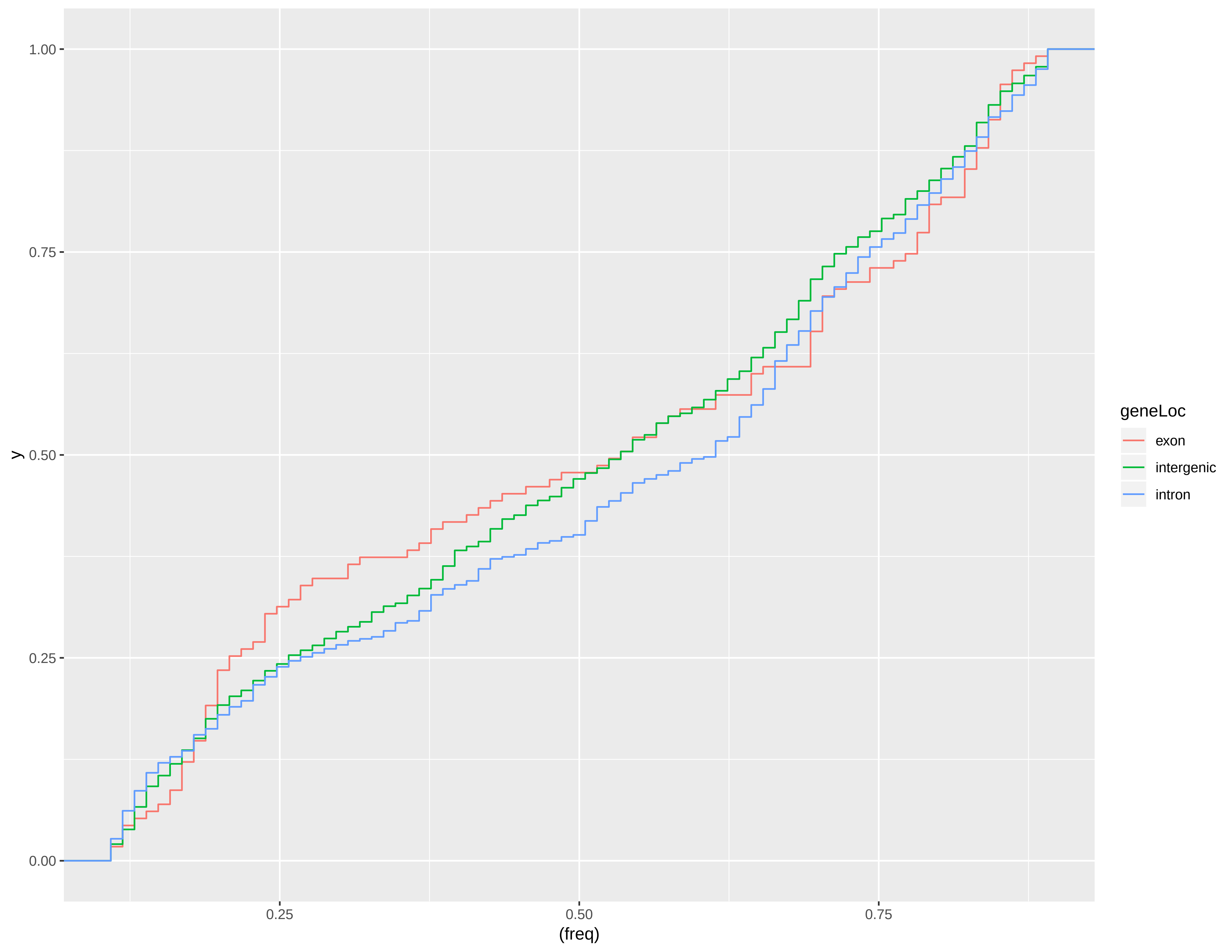
