## Supplemental Figure 3 for "Gene disruption by structural mutations drives selection in US rice breeding over the last century"

Welch Two Sample t−test

data: d$V6 by d$V7
t = −20.486, df = 11813, p−value < 2.2e−16
alternative hypothesis: true difference in means is not equal to 0
95 percent confidence interval:
−0.1461128 −0.1205936
sample estimates:
mean in group neutral mean in group selected
0.1941768 0.3275300

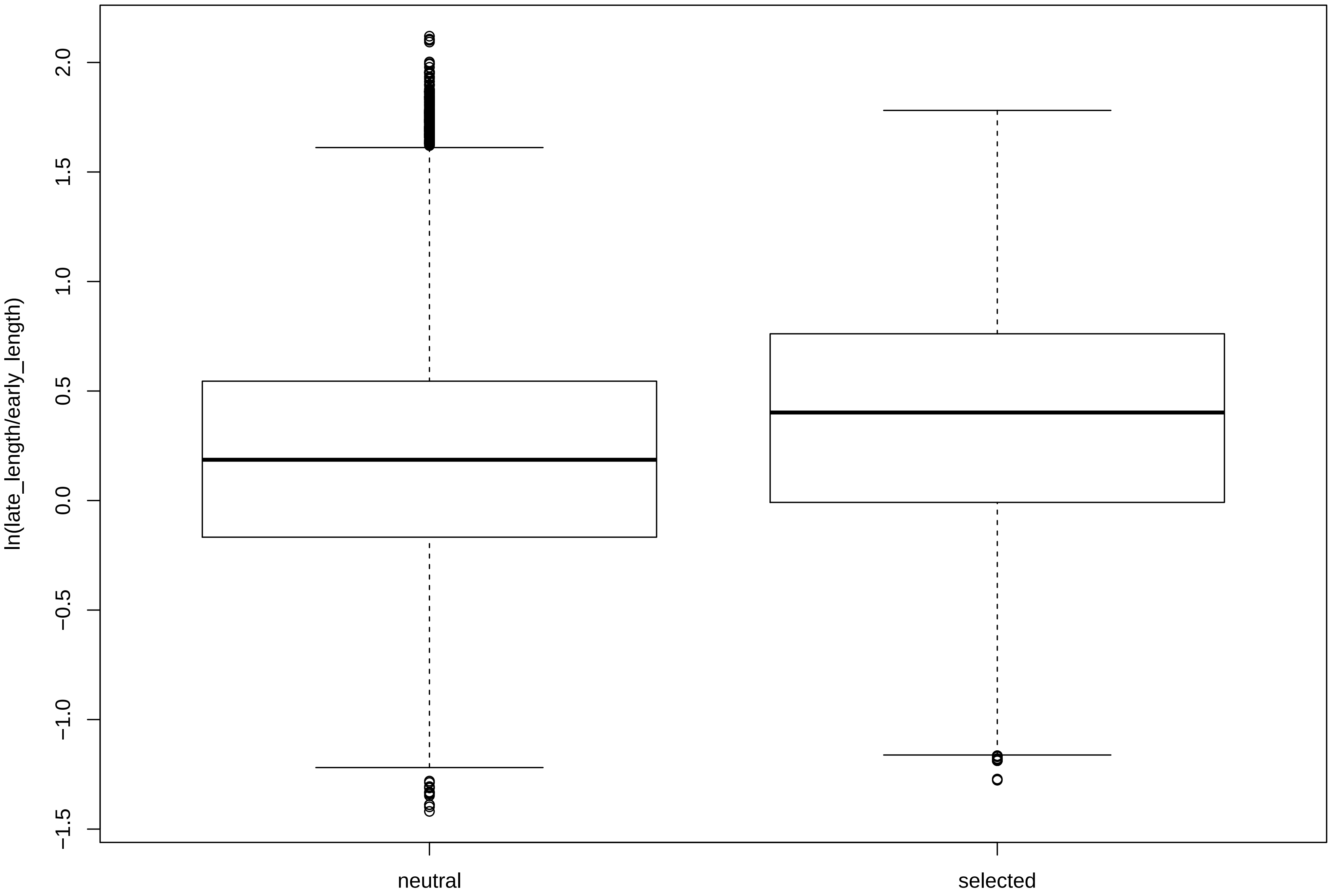
