## Supplemental Figure 4 for "Gene disruption by structural mutations drives selection in US rice breeding over the last century"

Post-filtering number of alignments: 21034□□□□minimum alignment length (-m): 10000
Post-filtering number of queries: 171□□□□□□□□minimum query aggregate alignment length (-q): 20000

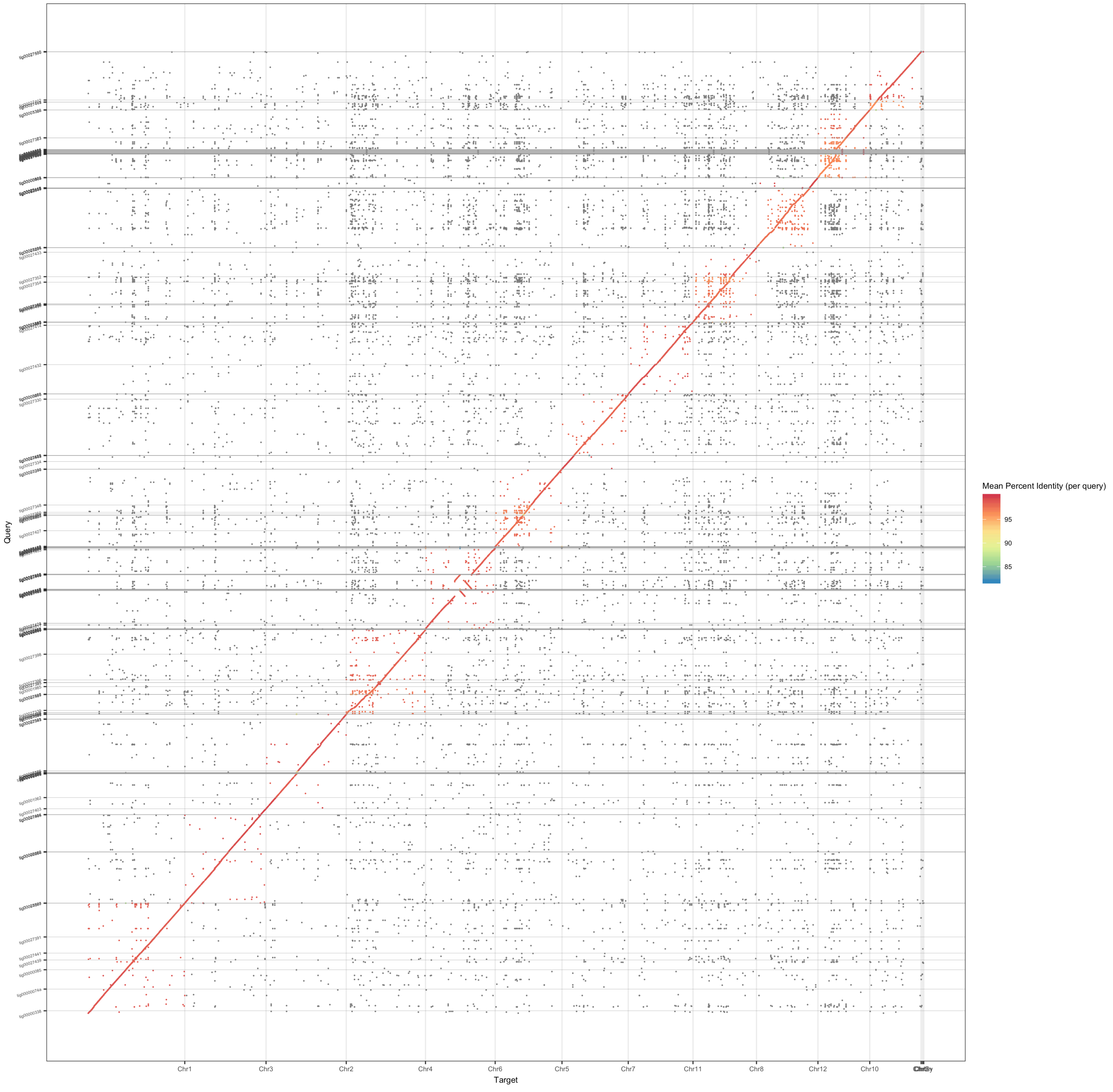
